## Supplemental material for "The Consortium for Genomic Diversity, Ancestry, and Health in Colombia (CÓDIGO): building local capacity in genomics, bioinformatics, and precision medicine"

Supplementary Figure 1. **Variant merging and harmonization.**

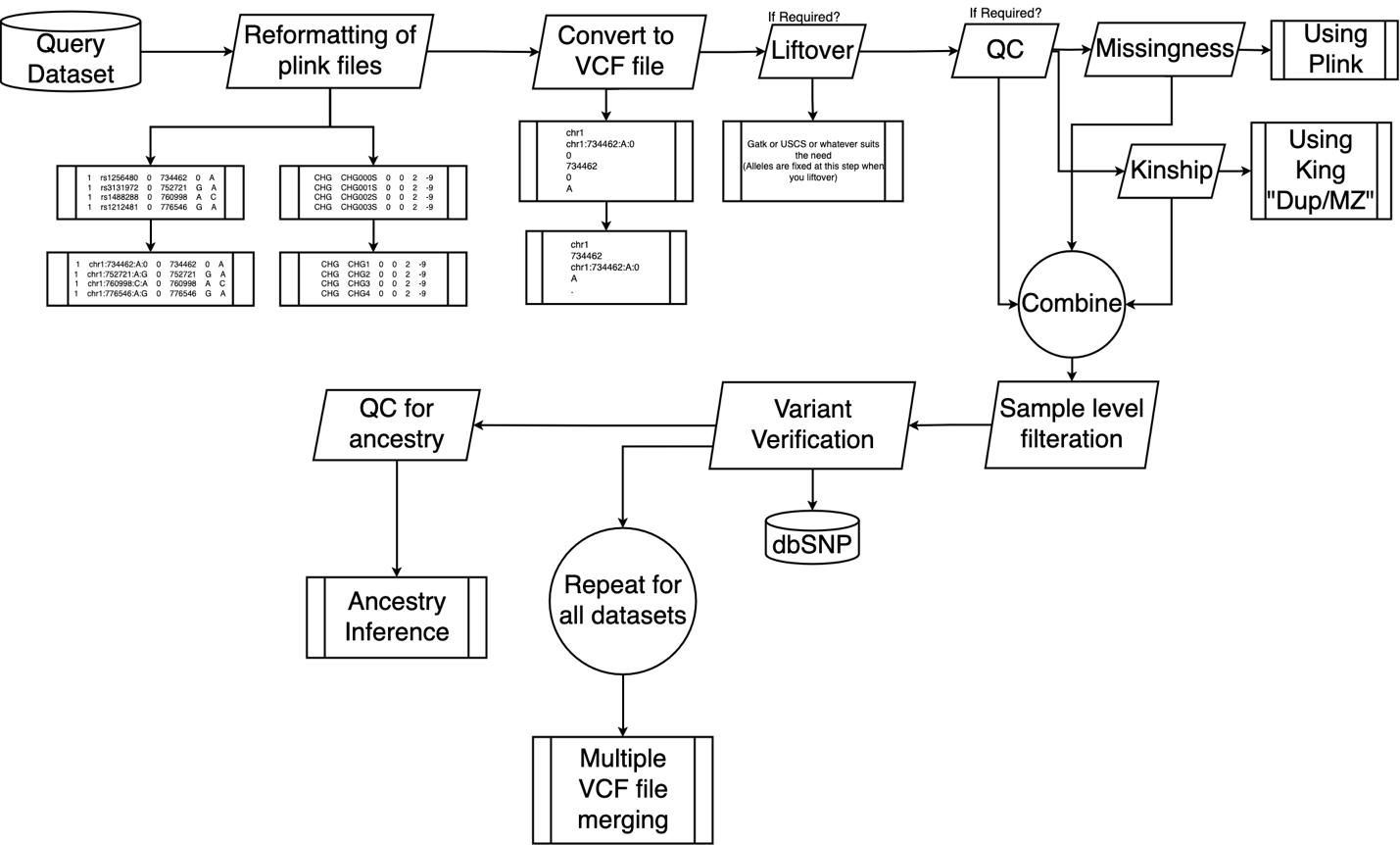

Supplementary Table 1. **CÓDIGO datasets.** The numbers (n) of samples and genomic variants before and after variant merging and harmonization are shown for each source dataset.

| **Source code^a^** | **Source description^b^** | **n samples before** | **n variants before** | **n samples after** | **n variants after** |
| --- | --- | --- | --- | --- | --- |
| CHG | Afro-Colombian from Chocó | 100 | 568,662 | 100 | 567,184 |
| PLQ | Afro-Colombian from San Basilio de Palenque | 34 | 10,064,050 | 34 | 9,779,781 |
| IND | Indigenous Arhuaco, Curripaco, Emberá, Guahibo, Inga, Kogi, Piapoco, Waunana, Wayuu communities | 50 | 364,430 | 50 | 296,141 |
| SIN | Indigenous Sinú community | 19 | 1,407,123 | 19 | 1,459,520 |
| CLM | Mestizo Colombian from Medellín | 94 | 81,568,731 | 94 | 81,568,727 |
| MCM | Mestizo Colombian from Medellín | 524 | 24,181,263 | 373 | 23,188,190 |
| MCA | Mestizo Colombian from Antioquia | 624 | 541,134 | 623 | 526,935 |
| VDC | Mestizo Colombian from Valle del Cauca | 116 | 6,481,171 | 116 | 6,481,171 |

^a^ Three letter code for each source dataset

^b^ Ethnicity and geographic origins for each source data set

Supplementary Table 2. **Global reference populations.** African, American, and European reference populations and samples used for genetic ancestry inference. Population descriptions taken from The International Genome Sample Resource (ISGR): <https://www.internationalgenome.org/data-portal/population>. 1KGP = 1000 Genomes Project; HGDP = Human Genome Diversity Project.

| **Name** | **Description** | **Superpopulation** | **n** | **Source** |
| --- | --- | --- | --- | --- |
| Esan | Esan in Nigeria | African | 99 | 1KGP |
| Gambian Mandinka | Gambian in Western Division, The Gambia - Mandinka | African | 106 | 1KGP |
| Yoruba | Yoruba in Ibadan, Nigeria | African | 107 | 1KGP |
| Karitiana | Karitiana in Brazil | American | 12 | HGDP |
| Maya | Maya in Mexico | American | 13 | HGDP |
| Peruvian | Peruvian in Lima, Peru | Admixed American | 14 | 1KGP |
| Pima | Pima in Mexico | American | 12 | HGDP |
| Surui | Surui in Brazil | American | 8 | HGDP |
| British | British in England and Scotland | European | 87 | 1KGP |
| Iberian | Iberian populations in Spain | European | 86 | 1KGP |
| Toscani | Toscani in Italy | European | 100 | 1KGP |

Supplementary Figure 2. **CÓDIGO development stack.** Server-side components used for the CÓDIGO webserver are shown.

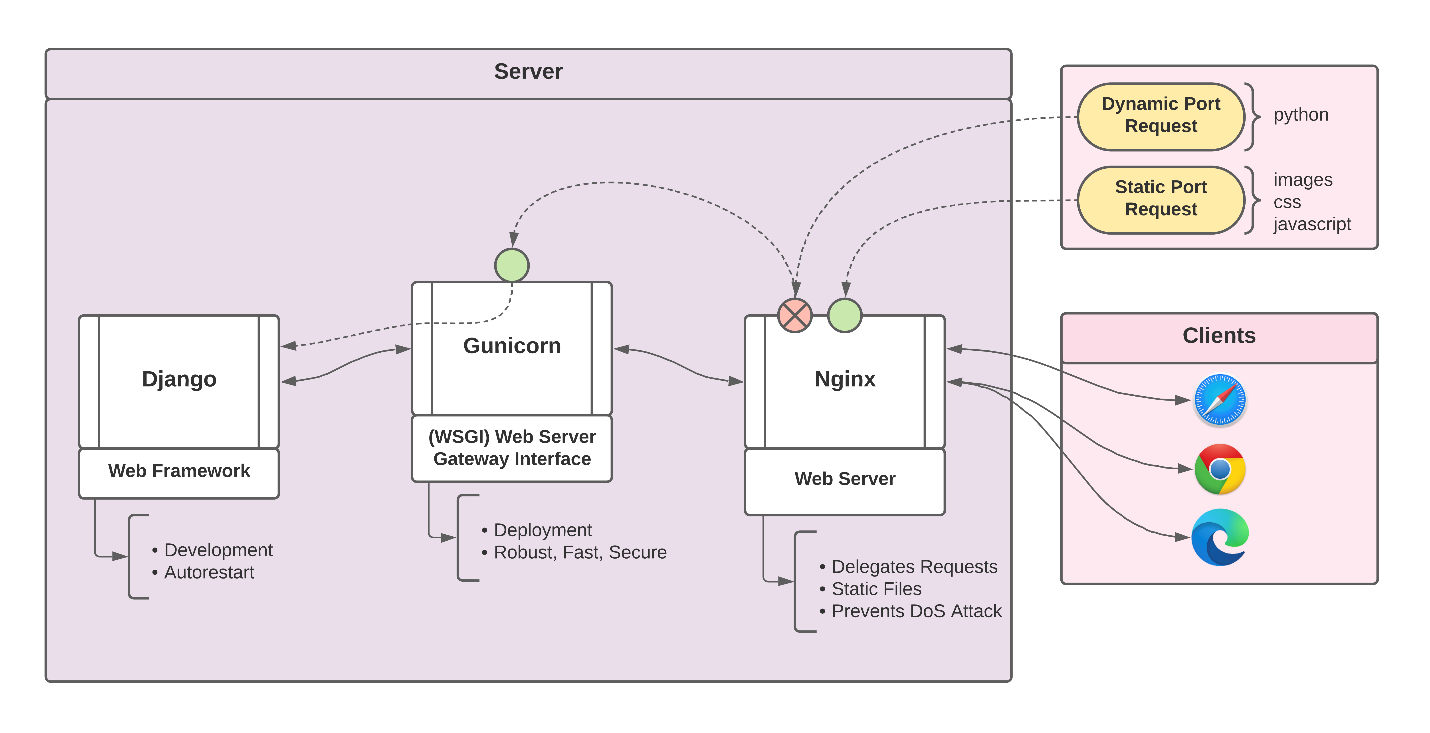

Supplementary Figure 3. **K-means clustering elbow plot.** Sum of squared errors (y-axis) plotted against the number of clusters (k; x-axis) for K-means clustering of CÓDIGO sample genetic ancestry fractions.

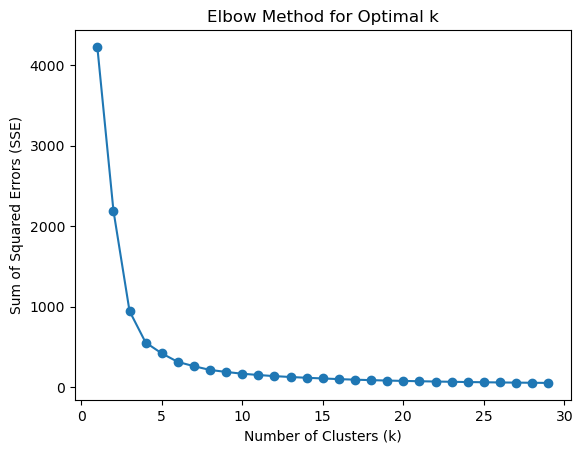
